# The flavonoid nobiletin exhibits differential effects on cell viability in keratinocytes exposed to UVA versus UVB radiation

**DOI:** 10.1101/2022.01.21.476505

**Authors:** William Cvammen, Michael G. Kemp

**Author notes:** Corresponding author (Michael G. Kemp).

## Abstract

The polymethoxylated flavonoid nobiletin has been shown to suppress inflammatory responses to UVB radiation and to enhance circadian rhythms. Because expression of the core nucleotide excision repair (NER) factor XPA and the rate of removal of UV photoproducts from DNA are regulated by the circadian clock, we investigated whether the beneficial effects of nobiletin in UVB-exposed cells could be due in part to enhanced NER. Though nobiletin limited UVB-irradiated human keratinocytes from undergoing cell death, we found that this enhanced survival was not associated with increased NER or XPA expression. Instead, nobiletin reduced initial UV photoproduct formation and promoted a G1 cell cycle arrest. We then examined the implications of this findings for exposures to solar radiation through use of a solar simulated light (SSL) source that contains primarily UVA radiation. In striking contrast to the results obtained with UVB radiation, nobiletin instead sensitized keratinocytes to both the SSL and a more defined UVA light source. This enhanced cell death was correlated with a photochemical change in nobiletin absorption spectrum in the UVA range. We conclude that nobiletin is unlikely to be a useful compound for protecting keratinocytes against the harmful effects of solar UV radiation.

## INTRODUCTION

Ultraviolet-B (UVB) wavelengths of sunlight induce the formation of potentially mutagenic photoproducts in the DNA of epidermal skin cells and are associated with the development of both melanoma and non-melanoma skin cancers. Natural products, synthesized chemicals, and other agents that either absorb this radiation or enhance UV photoproduct removal therefore have the potential to be used in sunscreens to limit photocarcinogenesis and photoaging (1, 2). Thus, there is significant interest in identifying compounds that can be added to sunscreens to increase their effectiveness and reduce the risk of skin cancer development.

One such possible compound is the citrus peel-derived, polymethoxylated flavonoid nobiletin (NOB), which has been shown to reduce inflammatory responses in human keratinocytes in vitro and mouse skin in vivo following exposure to UVB radiation (3). In addition, *C. elegans* treated with NOB exhibit enhanced resistance to UVC radiation and other stressors (4). Additional studies have found NOB to have antioxidant effects and to inhibit skin tumor formation in mice exposed to dimethylbenz[a]anthracene (5) or a nitric oxide donor (6). Thus, these results suggest that the possibility that NOB could be useful in preventing skin photocarcinogenesis and photoaging associated with UVB exposures.

Interestingly, NOB was also recently found to enhance the output of the circadian transcription machinery (7, 8) via agonism of retinoic acid-related orphan receptors (ROR), which act on ROR elements in the promoter of the BMAL1 gene to increase gene expression (9–11). Many aspects of mammalian physiology and biochemistry are known to be under circadian control (12, 13), including the rate of removal of UV photoproducts from DNA by the nucleotide excision repair (NER) machinery (14, 15). These oscillations in NER activity have been shown to be correlated with the expression of the core NER protein XPA throughout the day (16, 17). Thus, the timing of UVB exposure during the day or night influences both acute erythema and long-term skin carcinogenesis in mice (17, 18). Studies with human subjects similarly suggest that erythema and XPA expression exhibit circadian rhythmicity (19, 20).

Because of NOB’s reported beneficial functions in response to UV radiation (3, 4) and its ability to modulate circadian rhythms (7, 8), we were therefore curious whether NOB could be used protect human keratinocytes from UV radiation via BMAL1-mediated enhancement of XPA expression and UV photoproduct removal. However, we find here that although NOB reduces UVB lethality, the mechanism appears to be independent of XPA, NER, and effects on BMAL1 expression. Moreover, the effect of NOB is specific to UVB radiation, as exposure of NOB-treated cells to UVA or solar simulated UV radiation induces phototoxicity and a photochemical change in NOB.

## MATERIALS AND METHODS

### Cell culture

HaCaT keratinocytes were maintained in DMEM containing 10% FBS, an additional 2 mM L-glutamine, and penicillin-streptomycin. Telomerase-immortalized human neonatal foreskin keratinocytes (N-TERTs) (21) were cultured in EpiLife medium containing human keratinocyte growth supplement (HKGS) (Thermo Fisher Scientific) and penicillin/streptomycin. Both cell lines were maintained in a 5% CO_2_ humidified incubator at 37°C and monitored periodically for mycoplasma contamination (Sigma Venor GeM Kit). Unless otherwise indicated, cells were pre-treated with vehicle (0.2% DMSO) or 50 μM NOB for 24 hr and then exposed to the indicated UV light source in either culture medium or Hank’s balanced salt solution (HBSS). The UVB light source used two UVP XX-15M bulbs from Analytik Jena. The solar simulated light (SSL) source used two UVA340 bulbs from Q-Lab Corporation. Cells were exposed to UVA radiation by placing plates of cells under an inverted UVP Benchtop UV transilluminator set to the UVA (365 nm) setting. Fluence rates were determined with an International Light Technologies radiometer and UVB (SED240) or UVA (ILT72CE) sensors calibrated at 290 and 360 nm, respectively. Spectral properties of the light sources can be found in **Supplementary Figure S1**.

### Assays of cell survival and cell cycle distribution

Acute cell survival was determined using methylthiazolyldiphenyl-tetrazolium bromide (MTT), which was added to cell culture medium at a final concentration of 0.25 mg/ml and incubated for 30 min before solubilization with DMSO and measurement of absorbance at 570 nm on a Synergy H1 spectrophotometer (Bio-Tek). Absorbance values for the UV-irradiated samples were normalized to the non-irradiated samples. Measurements of cell death involved addition of propidium iodide (25 μg/ml) to unfixed suspensions of trypsinized cells harvested 24 hr after UV exposure. The percentage of PI-positive (dead) cells was determined using an Accuri C6 flow cytometer. To determine cell cycle distribution, ethanol-fixed cells were stained with 25 μg/ml propidium iodide after RNase-treatment and analyzed for DNA content using an Accuri C6 flow cytometer.

### DNA immunoblotting

Cells were treated with vehicle or NOB for 24 hr and then exposed to 200 J/m^2^ of UVB radiation. DNA was purified from cells harvested at the indicated time points using a Mammalian Genomic DNA Miniprep Kit (Sigma). DNA was immobilized on nitrocellulose and analyzed by immunodot blotting with antibodies against cyclobutane pyrimidine dimers (CPDs; Cosmo Bio NM-DND-001) and single-stranded DNA (Millipore MAB3034).

### Protein immunoblotting

Cells were lysed in ice-cold Triton X-100 lysis buffer (20 mM Tris-HCl, pH 7.5, 150 mM NaCl, 1 mM EDTA, 1 mM EGTA, and 1% Triton X-100) supplemented with 1X HALT protease inhibitor cocktail and incubated on ice for 15-20 min with occasional vortexing. Lysed cells were centrifuged at maximum speed in a refrigerated microcentrifuge. Protein lysates were separated on 8% or 15% Tris-Glycine SDS gels and then transferred to a nitrocellulose membrane using a semi-dry transfer apparatus. Blots were stained with 0.5% Ponceau S (Sigma) to ensure equal loading. The blots were blocked in in 5% non-fat milk in TBST (Tris-buffered saline containing 0.1% Tween-20) and then probed overnight with primary antibodies recognizing XPA, p21, BMAL1 or phosphorylated forms of KAP1, CHK1, CHK2, or histone H2AX. After washing with TBST, blots were probed with HRP-coupled anti-mouse or anti-rabbit IgG (Invitrogen) secondary antibodies for one hour at room temperature. Chemiluminescence was visualized with either Clarity Western ECL substrate (Bio-Rad) substrate using a Molecular Imager Chemi-Doc XRS+ imaging system (Bio-Rad). Signals in the linear range of detection were quantified by densitometry using Image Lab (Bio-Rad) and normalized to the Ponceau S-stained membranes.

### Nobiletin absorbance spectra

The absorbance spectra of nobiletin were determined using a NanoDrop One spectrophotometer after a 5 hr incubation in either the dark or under the solar simulated light source.

## RESULTS

### Nobiletin limits cells death in UVB-irradiated human keratinocytes

To determine whether NOB can protect keratinocytes from the lethal and/or anti-proliferative effects of UVB radiation, HaCaT keratinocytes were treated with NOB for 24 hr prior to exposure to increasing fluences of UVB radiation. As shown in **Figure 1a**, NOB-treated cells exhibited higher levels of cell survival as measured using an MTT assay. Similar results were seen in telomerase-immortalized, diploid neonatal (N-TERT) keratinocytes (**Figure 1b**). Dose-response experiments demonstrated that concentrations of NOB at 50 or 100 μM were required to promote UVB survival (**Supplementary Figure S2**). To determine whether this enhanced proliferation involved changes in cell death, vehicle- and NOB-treated HaCaT and N-TERT cells were analyzed for changes in cell membrane permeability by staining of unfixed cells with propidium iodide and analysis by flow cytometry 24 hr after UVB exposure. The results shown in **Figure 1c, d** demonstrated that NOB treatment resulted in reduced cell death in both cell lines. Clonogenic survival assays further demonstrated a protective effect of NOB on long-term proliferative potential in UVB-irradiated HaCaT cells (**Figure 1e**).

**Figure 1.**
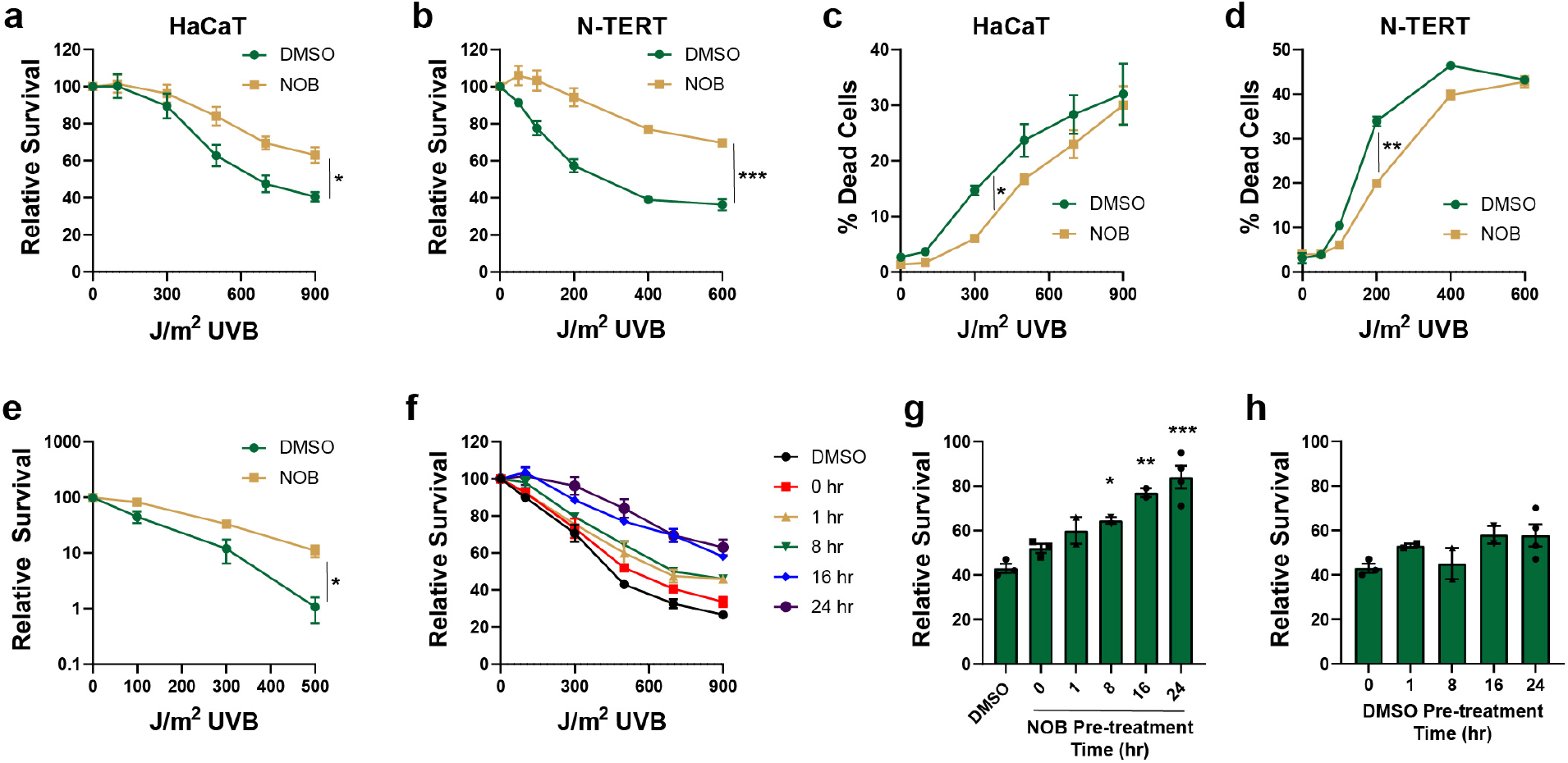
Nobiletin promotes keratinocyte survival in response to UVB radiation. **(a)** HaCaT cells were pre-treated with 50 μM nobiletin (NOB) for 24 hr before exposure to the indicated fluences of UVB radiation. MTT assays were performed 3 days later. **(b)** MTT assays were repeated as in (a) except with telomerase-immortalized diploid keratinocytes (N-TERTs) and pre-treatment with 100 μM NOB. **(c)** Cell death was measured by uptake of propidium iodide (PI) in non-fixed HaCaT treated as in (a) but harvested by trypsinization 24 after UVB exposure. **(d)** PI uptake was measured as in (c) except in N-TERT cells. **(e)** Clonogenic cell survival assays were performed in HaCaT cell treated as in (a). Cells were stained with crystal violet 2 weeks after UVB exposure and then quantified. **(f)** HaCaT cells were treated with 50 μM NOB for the indicated period of time before UVB exposure. MTT assays were performed 3 days later. **(g)** The relative level of cell survival at the 500 J/m^2^ UVB dose in (f) was graphed to show that NOB treatment exhibits a time-dependent promotion of UVB survival. **(h)** Data show the relative level of survival in cells treated with 0.2% DMSO vehicle in experiments performed as in (f). All data represent results from at least 3 independent experiments and analysis by one- or two-way ANOVA (*, p<0.05; **, p<0.01; ***, p<0.001).

If NOB acts in part via agonism of ROR and enhancement of the circadian transcriptional machinery, then the timing of NOB treatment prior to UVB exposure might be an important determinant of its effectiveness in mitigating the lethal effects of UVB radiation. Consistent with this hypothesis, a clear treatment time-dependent effect was observed in UVB-irradiated HaCaT cells, such that longer periods of pre-treatment with NOB (16 or 24 hr) resulted in greater cell survival than shorter pre-treatment times (**Figure 1f, g**). Importantly, this time-dependent effect was not observed with the DMSO vehicle (**Figure 1h**).

### Nobiletin does not enhance XPA expression or UV photoproduct removal rate but reduces initial UV photoproduct formation and DNA damage response kinase signaling

Because UV photoproduct removal by NER and expression of the NER protein XPA are controlled by the circadian clock transcriptional machinery (16, 17, 22–24), the activity of which is stimulated by NOB (7, 8), we next examined whether the pro-survival effect of NOB was mediated in part via enhanced XPA expression and/or NER rate. However, as shown in **Figure 2a-b**, treatment of HaCaT cells for 24 hr with NOB did not lead to significant changes in XPA protein expression. Moreover, NOB treatment instead reduced XPA expression in N-TERT keratinocytes (**Figure 2c**).

**Figure 2.**
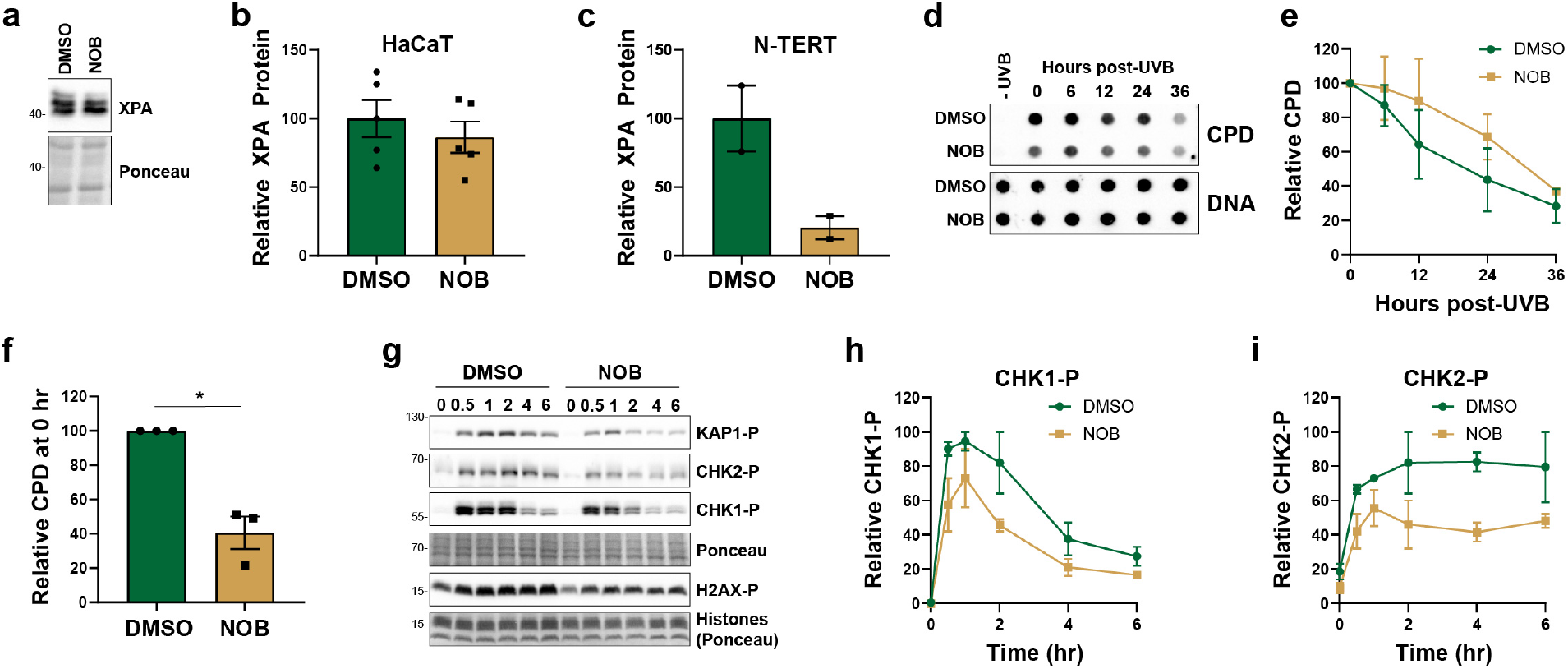
Nobiletin does not promote XPA expression or nucleotide excision repair activity but limits initial UV photoproduct formation. **(a)** Representative western blot for XPA protein in HaCaT cells treated with 50 μM NOB or 0.2% DMSO vehicle for 24 hr. **(b)** Quantitation of XPA expression in 5 independent experiments performed as in (a). **(c)** Quantitation of XPA expression in two independent experiments performed as in (a) but with N-TERT keratinocytes. **(d)** Representative DNA immunoblot showing cyclobutane pyrimidine dimer (CPD) formation and repair after exposure of cells treated as in (a) to 200 J/m^2^ UVB. **(e)** Quantitation of CPD removal from genomic DNA from 3 independent experiments performed as in (d). **(f)** Quantitation of initial CPD formation from experiments in (d). Relative CPD content was analyzed by one-sample t-test. **(g)** HaCaT cells were treated as in (a), exposed to 200 J/m^2^ UVB, and then harvested at the indicated time points for western blot analysis. **(h, i)** Quantitation of CHK1 and CHK2 phosphorylation from 2 independent experiments performed as in (g).

To determine whether the activity of the NER system is affected by NOB treatment, we next analyzed the removal of cyclobutane pyrimidine dimers (CPDs) from DNA in UVB-irradiated HaCaT cells by DNA immunoblotting. As shown in **Figure 2d-e**, NOB treatment had no significant effect on the rate of CPD removal from genomic DNA. However, we noted in our DNA immunoblots in **Figure 2d** that the level of CPD formation at time 0 (immediately after UVB exposure) was reduced in cells treated with NOB. Quantitation of CPD formation at this time point from three independent experiments showed that NOB reduced the initial amount of CPDs that formed in genomic DNA by greater than 50% (**Figure 2f**). Consistent with reduced CPD formation in NOB-treated cells exposed to UVB radiation, DNA damage response kinase signaling was partially reduced in NOB-treated cells (**Figure 2g**), which included phosphorylation of the checkpoint kinases CHK1 and CHK2 (**Figure 2h-i**). Thus, we conclude from these experiments that NOB treatment leads to reduced CPD formation after UVB exposure but does not alter the activity of the NER machinery or expression of its key clock-regulated protein XPA.

### Nobiletin treatment leads to an increased number of cells in the G1 phase of the cell cycle in a p21-independent manner

Progression through the cell cycle has also been shown to be under circadian control (25, 26). We therefore next examined whether NOB altered the distribution of cells throughout the cell cycle by flow cytometry. As shown in **Figure 3a**, treatment of HaCaT cells with NOB for 24 hr was associated with increased numbers of cells in the G1 phase of the cell cycle and a reduced fraction in S and G2 phase. A similar, statistically significant reduction in S phase cells was observed in NOB-treated N-TERT cells (**Figure 3b**). Prolonged treatment of these cells with high concentrations of NOB led to a noticeable reduction in cell proliferation without any dramatic changes in cell viability (**Supplementary Figure S3**).

**Figure 3.**
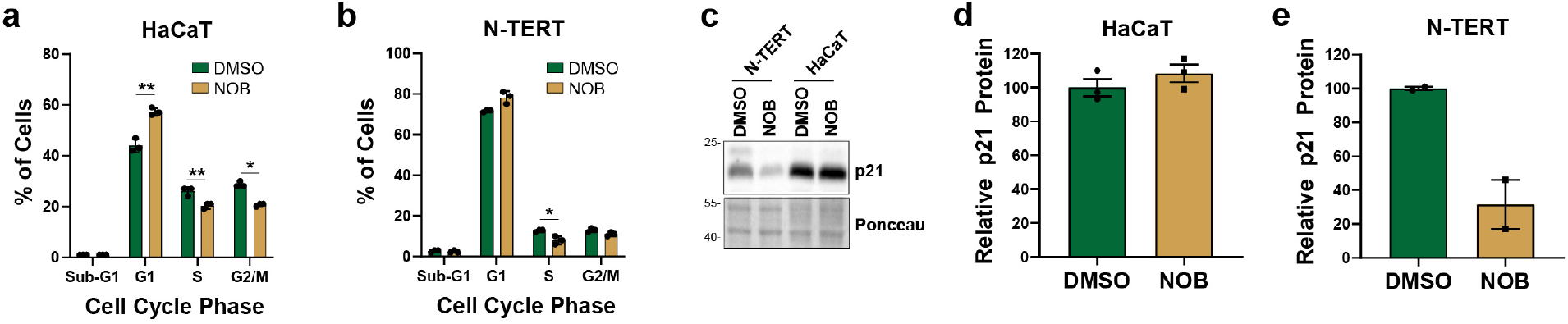
NOB alters cell cycle distribution independent of the cyclin-dependent kinase inhibitor protein p21. **(a)** HaCaT keratinocytes were treated with vehicle (DMSO) or NOB for 24 hr. Cell cycle distribution was determined by staining of fixed cells with propidium iodide and analysis by flow cytometry. **(b)** N-TERT keratinocytes were treated as in (a). Paired t-tests were used to determine whether NOB induced a significant change in cell cycle distribution (*, p<0.05; **, p<0.01). **(c)** Representative western blot for p21 protein in N-TERT and HaCaT keratinocytes treated with DMSO or NOB for 24 hr. **(d-e)** Quantitation of p21 protein levels from HaCaT and N-TERTs cells from 2-3 experiments performed as in (c).

The cyclin-dependent kinase inhibitory protein p21 (27), which is known to promote arrest in G1 of the cell cycle, has been shown to be regulated by BMAL1 (25). However, we found that NOB treatment did not enhance p21 expression in either HaCaT or N-TERT keratinocytes (**Figure 3c-e**). Because cells in S phase are more sensitive to UV radiation (28), we suggest that the enrichment of cells in G1 by NOB treatment may contribute to the resistance of cells to UVB radiation. However, this effect is likely not mediated by the previously reported regulation of p21 expression (25).

### Nobiletin does not enhance BMAL1 protein expression

Because the RORs transcriptionally activate BMAL1 expression (9–11), the increased UVB survival seen with the ROR agonist NOB treatment might be due promoting BMAL1 expression. However, the results presented thus far indicated to us that NOB may be acting to protect keratinocytes from UVB radiation via a mechanism that is independent of the known clock control genes XPA and p21. Consistent with this latter hypothesis, treatment of keratinocytes with NOB for 24 hr led to only a very modest 8% increase in BMAL1 protein levels in HaCaT cells (**Figure 4a, b**) and instead led to a 40% decrease in N-TERT cells (**Figure 4a, c**). Thus, we conclude that NOB promotes keratinocyte survival in response to UVB radiation by a mechanism that is likely independent of the canonical circadian transcription machinery. Instead, reduced CPD formation and enhanced G1 cell cycle arrest by NOB may be responsible for the pro-survival effect of NOB in response to UVB radiation.

**Figure 4.**
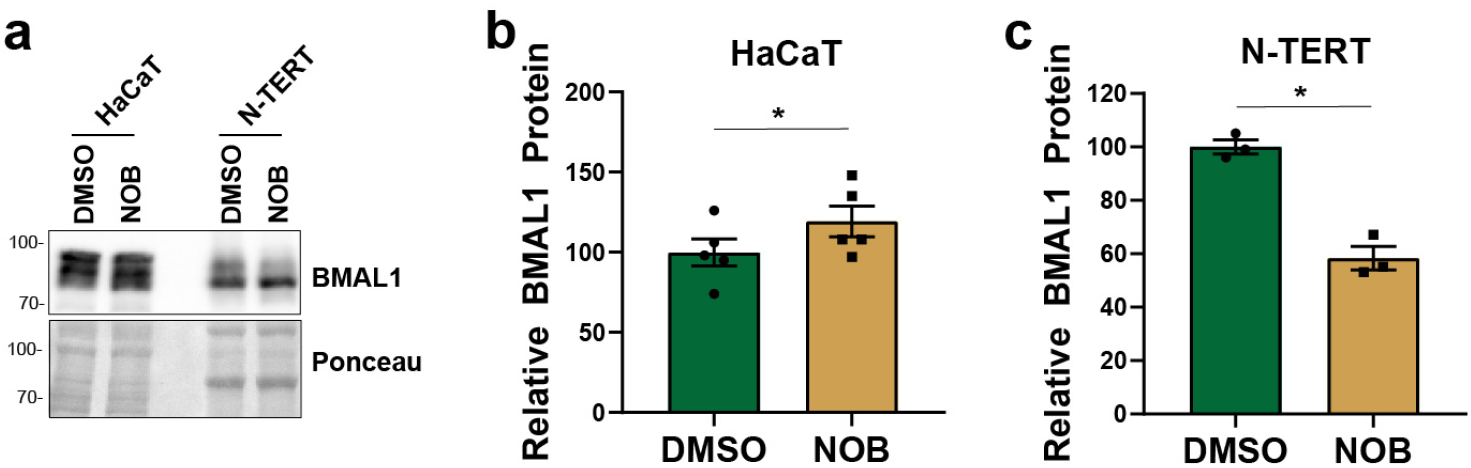
Nobiletin does not enhance BMAL1 protein expression in human keratinocytes. **(a)** Representative western blot for BMAL1 protein in HaCaT and N-TERT cells treated with DMSO or NOB for 24 hr. **(b)** Quantitation of BMAL1 expression in HaCaT cells from 5 independent experiments performed as in (a). **(c)** Quantitation of BMAL1 expression in three independent experiments performed as in (a) but with N-TERT keratinocytes. Ratio t-tests were used to compare relative BMAL1 protein levels.

### Nobiletin sensitizes keratinocytes to UVA radiation

Though our data indicate NOB promotes UVB survival through a mechanism inconsistent of enhanced DNA repair, our results nonetheless suggest that NOB could be useful in mitigating some of the negative consequences of UVB exposure. However, the UVB light source used in our experiments above does not accurately represent the distribution of solar UV wavelengths that humans and other terrestrial organisms are routinely exposed to. We therefore repeated our cell survival assays using a solar simulated light (SSL) source that is composed of approximately 93% UVA and 7% UVB (29). Interestingly, as shown in **Figure 5a**, pre-treatment of HaCaT cells with NOB for 24 hr failed to protect the cells from SSL and instead greatly sensitized the cells to the anti-proliferative effects of this light source. Moreover, we noticed that pre-treatment of the cells was not necessary for this sensitization, as treatment of cells with NOB in either culture media or Hank’s balanced salt solution (HBSS) solely during the period of SSL exposure (and not before or after SSL exposure) led to similar levels of SSL sensitivity (**Figure 5b, c**). Furthermore, treatment of cells with NOB after SSL exposure did not significantly sensitize the cells to SSL (**Figure 5d**). Finally, assays measuring cell death via uptake of propidium idodide in non-fixed cells confirmed that NOB induces phototoxicity in response to SSL exposure in both HaCaT (**Figure 5e**) and N-TERT (**Supplementary Figure S4**) keratinocytes.

**Figure 5.**
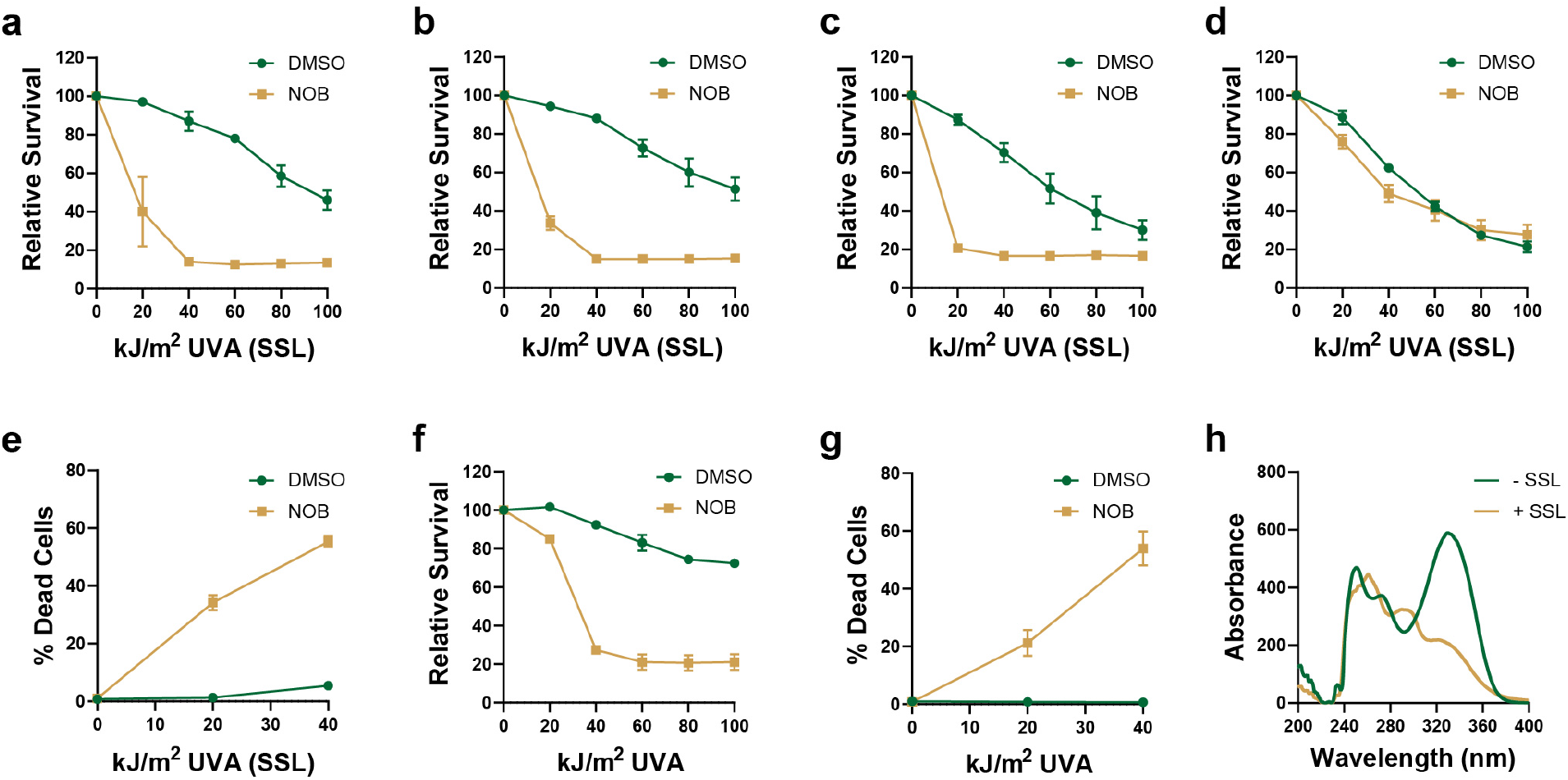
Nobiletin sensitizes keratinocytes to UVA radiation. **(a)** HaCaT cells were pre-treated with 50 μM nobiletin (NOB) for 24 hr before exposure to the indicated fluences of UVA radiation in a solar simulated light (SSL) source. MTT assays were performed 3 days later. **(b)** MTT assays were repeated as in (a) except that DMSO or NOB was added to cell culture medium immediately prior to SSL exposure and then washed at the end of the exposure period. **(c)** Cells were treated as in (b) except that DMSO/NOB was added to Hank’s buffered salt solution (HBSS) instead of cell culture media. **(d)** Cells were treated as in (c) except that NOB was added to cell culture media after the SSL exposure. **(e)** Cell death assays were performed by measuring propidium idodide uptake in non-fixed cells 24 hr after exposure of HaCaT cells SSL in HBSS containing DMSO or NOB. **(f)** HaCaT cells were treated as in (a) except cells were exposed to a UVA light source. **(g)** Cell death was measured as in (e) except with cells exposed to UVA instead of the SSL source. **(h)** The absorbance spectrum of NOB following a 5 hr exposure to the SSL source or incubation in the dark. With the exception of the data in (d), two-way ANOVAs showed that NOB-treated cells exhibited a significant difference in cell survival.

To further characterize the differential responses of NOB-treated cells to UVB radiation versus solar simulated light, we next exposed NOB-treated HaCaT cells to a UVA light source with less contaminating UVB radiation (**Supplementary Figure S1**). As shown in **Figure 5g-h**, NOB greatly sensitized the cells to UVA radiation, indicating that the UVA component of the SSL is responsible for the NOB photosensitization.

Our data showing that NOB limits CPD induction by UVB radiation and induces UVA photosensitization suggests that NOB may be absorbing UVA and UVB wavelengths of light. We therefore next determined the absorption spectrum of NOB in the absence and presence of prolonged exposure to SSL. As shown in **Figure 5h**, NOB displayed several absorption peaks within the UV range, including a major absorption peak at approximately 340 nm (within the UVA range). No absorption was observed at wavelengths above 400 nm (data not shown). Interestingly, we observed a time-dependent reduction in the 340 nm absorption peak after exposure to SSL, such that the peak was nearly completely gone after 5 hr exposure. These results indicate that NOB undergoes a photochemical change in response to UVA exposure and that this may be associated with the NOB phototoxicity in the presence of UVA wavelengths of light. Thus, we conclude that though NOB promotes survival in response to UVB exposure, it has a strikingly different (and instead toxic) effect in response to light sources containing significant levels of UVA radiation.

## DISCUSSION

There is great interest in improving the effectiveness of sunscreens to protect vulnerable individuals from mutagenesis and carcinogenesis associated with exposures to solar UV radiation, including through the use of natural products and the application of sunscreens containing DNA repair enzymes (1, 2, 30). Because the natural product NOB has been shown to enhance the transcriptional output of the circadian transcriptional machinery (7), which is known to regulate UV photoproduct removal and other UVB DNA damage responses (14, 15, 17), we investigated value of NOB in keratinocyte responses to UV radiation. Though NOB protected both HaCaT and telomerase-immortalized diploid keratinocytes from UVB radiation, we found this mechanism to be independent of DNA repair and the core circadian clock protein BMAL1. Because the effect of NOB on UVB survival was treatment time-dependent and is associated with decreased UVB photoproduct induction and altered cell cycle progression, there may be multiple mechanisms by which NOB protects keratinocytes from the lethal effects of UVB radiation.

Though UVB photoprotection is important, we unexpectedly found that NOB instead sensitizes keratinocytes to UV light sources composed primarily of UVA radiation, which more closely mimics the distribution of UV light in solar radiation. Our observation that NOB exhibits a loss in absorbance of NOB at 340 nm after exposure to SSL indicates that the compound undergoes a photochemical change in response to UVA radiation. The nature of this change and whether it is responsible for the UVA phototoxicity observed here remains to be determined. Nonetheless, the results suggest that NOB is not a useful compound for protecting keratinocytes from solar UV radiation. Thus, studies using UVB light sources along with NOB treatment should be reviewed with caution. However, the fact that NOB induces a cell cycle arrest could be useful to protect cells from the lethal and mutagenic effects of other DNA damaging agents and may contribute to the anti-tumor effects of NOB in the context of treatment with dimethylbenz[a]anthracene (5). Finally, it is also possible that the effects of NOB may differ in cultured cells that lack circadian synchronization versus cells or other organ systems in vivo that possess robust circadian rhythmicity.

## ACKNOWLEDGMENTS

The authors thank the WSU Proteome Analysis Laboratory and Center for Genomics Research for the use of equipment to carry out this work. This work was supported by start-up funding provided by Wright State University and by grants from the National Institute of General Medical Sciences (GM130583), Ohio Cancer Research Associates (#5020), and the Veterans Administration (I01CX002241).

