## Supplementary Figures for "The flavonoid nobiletin exhibits differential effects on cell viability in keratinocytes exposed to UVA versus UVB radiation"

**a**

| UV Fluence Rates (J/m²/sec) | | | | | | |
|---|---|---|---|---|---|---|
| | | | Vendor: | UVP | Q-Lab | UVP |
| | | | Light Source: | XX-15M (UVB) | UVA340 (SSL) | Trans (365 nm UVA) |
| Vendor Radiometer | Sensor | Peak Detection | Distance: | 12 in | 4 in | 3.5 in |
| UVP | UVX-25 | 254 nm | | 5.0 | 1.8 | 1.2 |
| ILT | SED240 | 290 nm | | 9.0 | 5.2 | 0.1 |
| UVP | UVX-31 | 310 nm | | 12.0 | 26.3 | 7.6 |
| ILT | ILT72CE | 360 nm | | 2.6 | 44.3 | 27.0 |
| UVP | UVX-36 | 365 nm | | 2.5 | 33.0 | 31.8 |

**b**

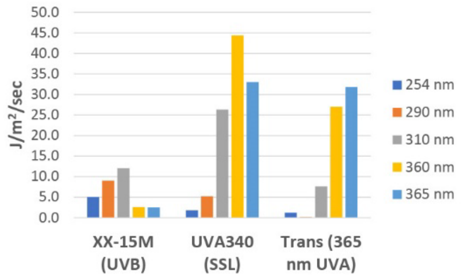


**Figure S1. Properties of UV light sources. (a)** Characteristics of the UV light sources used in this work. For each light source, the vendor and product, distance between the light source and cells, and the fluence rate at specific wavelengths were measured and are provided in the table. **(b)** Graphical representation of the fluence rate of UV at each of the indicated wavelengths for the three UV light sources used here.

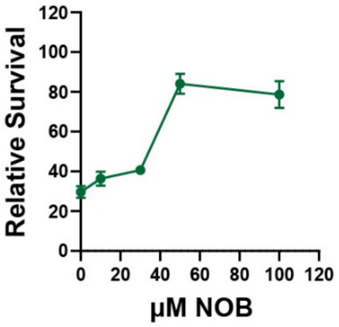


**Figure S2. High doses of NOB are required to protect HaCaT keratinocytes from UVB exposure.** HaCaT keratinocytes were exposed to the indicated dose of NOB before exposure to 500 J/m$^2$ UVB radiation. MTT assays were performed 3 days later. The graph shows the average level of survival from 3-4 independent experiments.

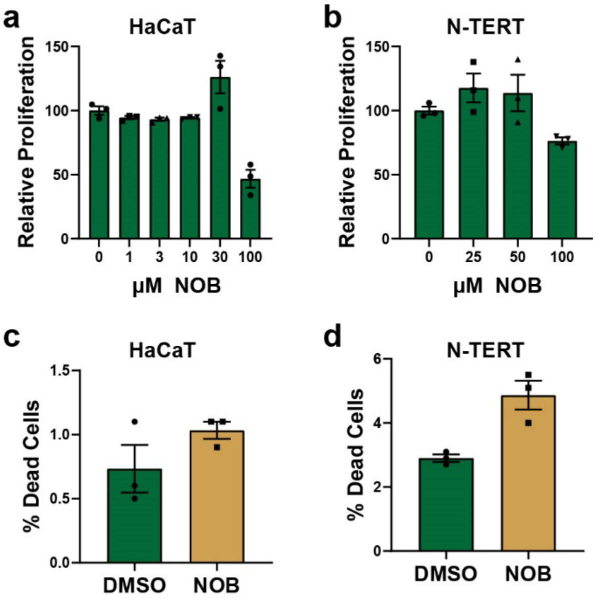


**Figure S3. High doses of NOB inhibit keratinocyte proliferation. (a)** HaCaT keratinocytes were exposed to the indicated dose of NOB for 3 days. MTT assays were to examine relative proliferation. **(b)** N-TERTs were treated as in (a). The graph shows the average level of relative proliferation from 3 independent experiments. **(c)** HaCaT cell viability was determined by PI uptake after 24 hr of treatment with 50 µM NOB. **(d)** N-TERT cell viability was determined 24 hr after treatment with 100 µM NOB.

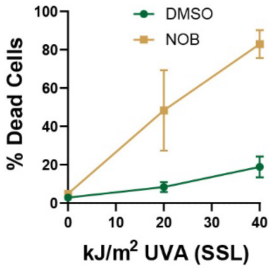


**Figure S4. NOB sensitizes N-TERT keratinocytes to solar simulated light.** N-TERTs were incubated in Hank's buffered salt solution containing DMSO or 100 µM NOB and then exposed to the indicated fluence of solar simulated light (SSL) as measured with a UVA radiometer at 360 nm.
